## Supplemental Figures, Sup Table 1, and Sup Table 2 for "G Protein-Coupled Receptor Kinase-2 (GRK-2) controls exploration through neuropeptide signaling in *Caenorhabditis elegans*"

#### Figure S1

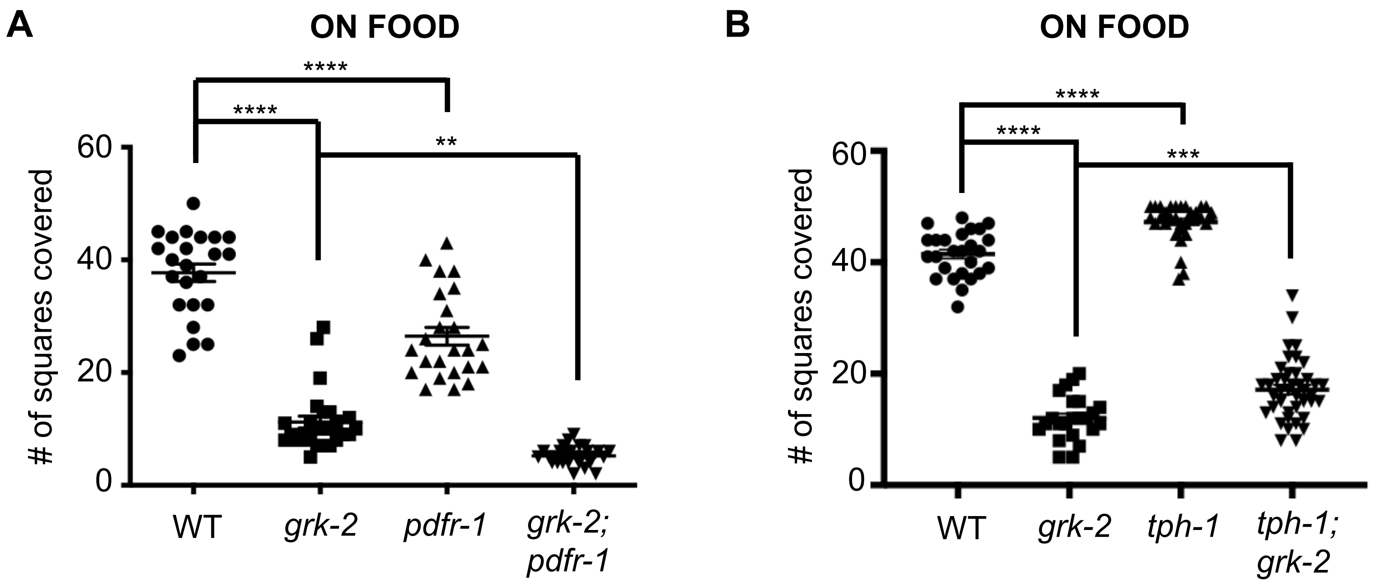
A
ON FOOD
****
****
**
60
40
20
0
### of squares covered
WT
grk-2
pdfr-1
grk-2;
pdfr-1
B
ON FOOD
****
****
***
60
40
20
0
### of squares covered
WT
grk-2
tph-1
tph-1;
grk-2

**Figure S2**

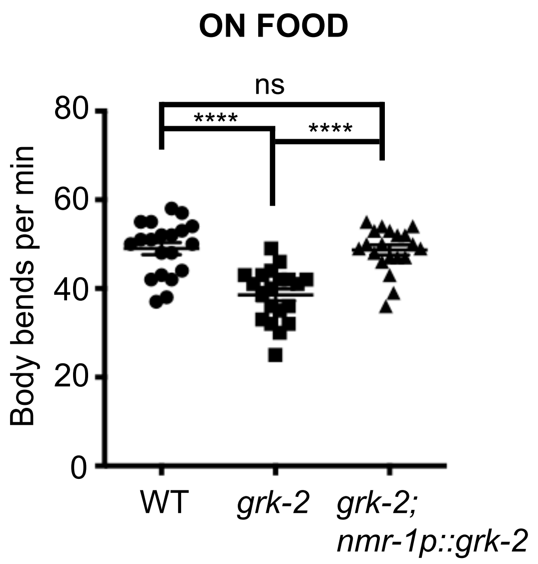
ON FOOD
ns
****
****
80
60
40
20
0
Body bends per min
WT
grk-2
grk-2;
nmr-1p::grk-2

**Figure S3**

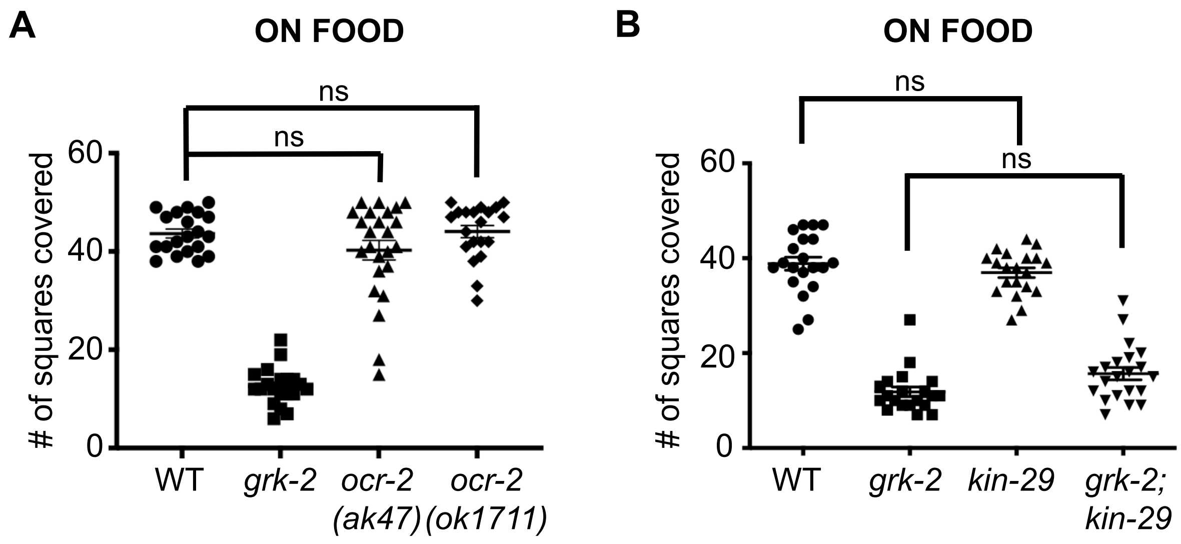
A
ON FOOD
ns
ns
### of squares covered
60
40
20
0
WT
grk-2
ocr-2 (ak47)
ocr-2 (ok1711)
B
ON FOOD
ns
ns
### of squares covered
60
40
20
0
WT
grk-2
kin-29
grk-2; kin-29

**Figure S4**

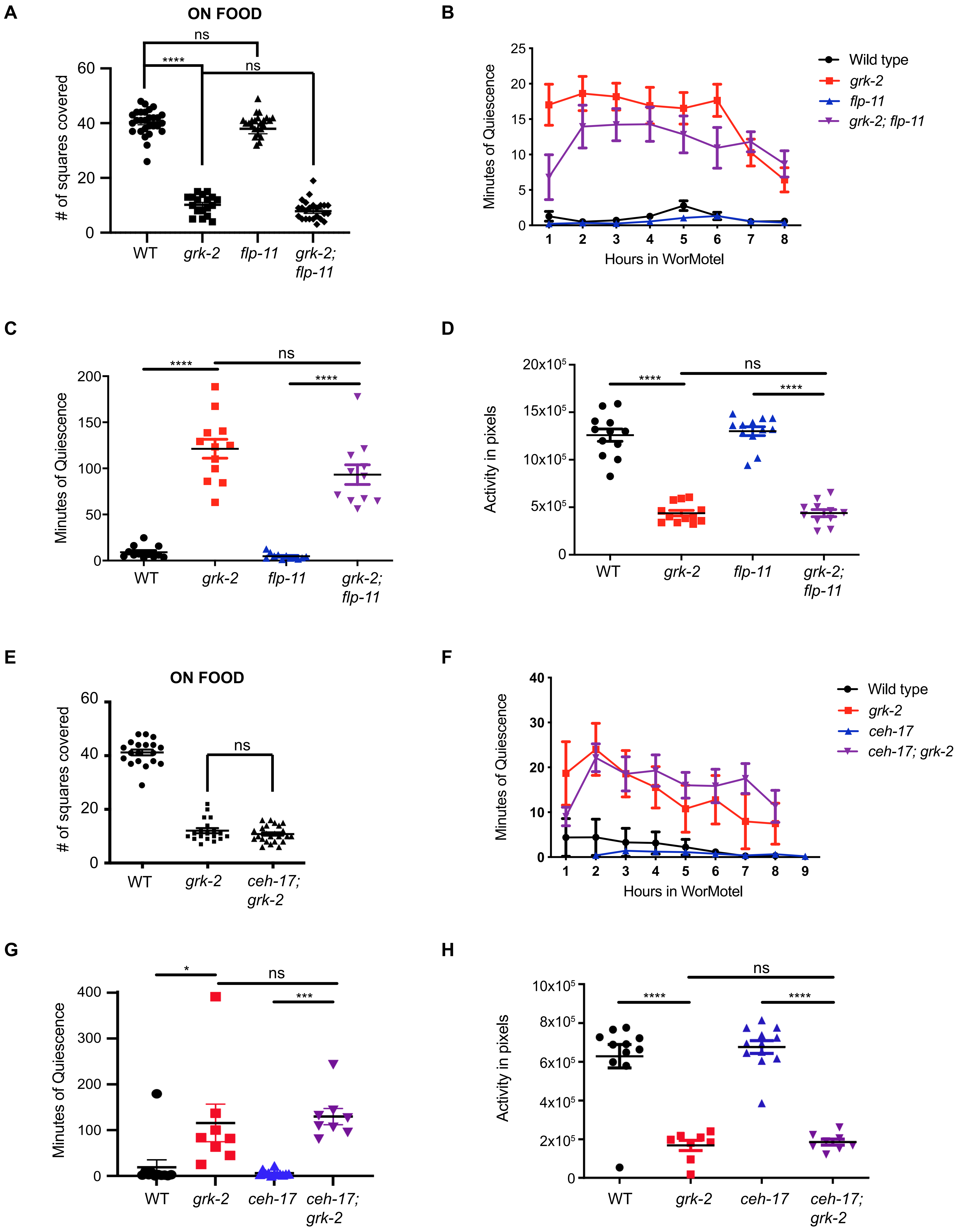
A
ON FOOD
ns
****
ns
### of squares covered
60
40
20
0
WT
grk-2
flp-11
grk-2; flp-11
B
Minutes of Quiescence
25
20
15
10
5
0
1 2 3 4 5 6 7 8
Hours in WorMotel
Wild type
grk-2
flp-11
grk-2; flp-11
C
****
ns
****
Minutes of Quiescence
200
150
100
50
0
WT
grk-2
flp-11
grk-2; flp-11
D
****
ns
****
Activity in pixels
20x10^5
15x10^5
10x10^5
5x10^5
0
WT
grk-2
flp-11
grk-2; flp-11
E
ON FOOD
ns
### of squares covered
60
40
20
0
WT
grk-2
ceh-17; grk-2
F
Minutes of Quiescence
40
30
20
10
0
1 2 3 4 5 6 7 8 9
Hours in WorMotel
Wild type
grk-2
ceh-17
ceh-17; grk-2
G
*
ns
***
Minutes of Quiescence
400
300
200
100
0
WT
grk-2
ceh-17
ceh-17; grk-2
H
ns
****
****
Activity in pixels
10x10^5
8x10^5
6x10^5
4x10^5
2x10^5
0
WT
grk-2
ceh-17
ceh-17; grk-2

**Figure S5**

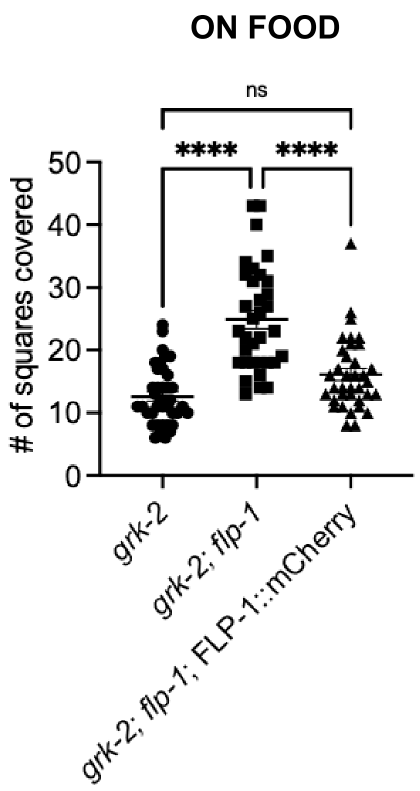

ON FOOD
ns
****
****
### of squares covered
50
40
30
20
10
0
grk-2
grk-2; flp-1
grk-2; flp-1; FLP-1::mCherry

**Supporting information**

**S1 Table. List of strains.**

N2: Bristol wild strain

AX1295: *gcy-35(ok769)* I

AX1410: *flp-18(db99)* X

CB1033: *che-2(e1033)* X

CB1126: *che-6(e1126)* IV

CB1377: *daf-6(e1377)* X

CB1338: *mec-3(e1338)* IV

CB1339: *mec-4(e1339)* X

CB1387: *daf-10(e1387)* IV

CB3323: *che-13(e1805)* I

CB3329: *che-10(e1809)* II

CB3330: *che-11(e1810)* V

CB3332: *che-12(e1812)* V

CB3687: *che-14(e1960)* I

CX10: *osm-9(ky10)* IV

CX32: *odr-10(ky32)* X

CX2065: *odr-1(n1936)* X

CX2304: *odr-2(n2145)* V

CX2357: *odr-5(ky9)* X

CX2386: *odr-8(ky31)* IV

CX3222: *odr-3(n1605)* V

CX4148: *npr-1(ky13)* X

FG7: *grk-2(gk268)* III

FX1497: *npr-6(tm1497)* X

HBR227: *aptf-1(gk794)* II

HBR507*: flp-11(tm2706)* X

IB16: *ceh-17(np1)* I

JT7641: *odr-7(ky4)* X

JT7674: *tax-4(p678)* III

LX703*: dop-3(vs106)* X

MT1074*: egl-4(n479)* IV

MT3564: *osm-7(n1515)* III

MT3641: *osm-10(n1602)* III

MT3643: *osm-11(n1604)* X

MT5300: *odr-4(n2144)* III

MT15434: *tph-1(mg280)* II

OH13098: *che-1(ot75)* I

PR691: *tax-2(p691)* I

PR813: *osm-5(p813)* X

PY1479: *kin-29(oy38)* X

RB982: *flp-21(ok889)* V

RB1834: *che-7(e1128)* V

VC1233: *ocr-2(ok1711)* IV

VC2609: *pdfr-1(ok3425)* III

VM396: *ocr-2(ak47)* IV

VC40103: *frpr-7(gk463846)* X

XZ1544: *grk-2(gk268)* III ; *yakEx44[rab-3p::grk-2 cDNA::tbb-2* 3'UTR::OPERON::GFP*, myo-3p::mCherry]*

XZ1551: *grk-2(gk268)* III ; *yakEx47[acr-2p::grk-2 cDNA::tbb-2* 3'UTR::OPERON::GFP*, myo-2p::mCherry]*

XZ1903*: grk-2(gk268)* III ; *dop-3(vs106)* X

The following strains were produced in this study:

BJH1124: *grk-2(gk268)* III *; pekEx265[str-1p::grk-2 cDNA::tbb-2* 3'UTR::OPERON::GFP*, unc-122*::GFP*]*

BJH1125: *grk-2(gk268)* III *; pekEx265[str-2p::grk-2 cDNA::tbb-2* 3'UTR::OPERON::GFP*, unc-122*::GFP*]*

BJH1126: *grk-2(gk268)* III *; pekEx266[odr-10p::grk-2 cDNA::tbb-2* 3'UTR::OPERON::GFP*, unc-122*::GFP*]*

BJH2357: *grk-2(gk268)* III ; *flp-21(ok889)* V

BJH2361: *grk-2(gk268)* III ; *daf-10(e1387)* IV

BJH2371: *grk-2(gk268)* III ; *flp-1(ok2811)* IV

BJH2762: *ocr-2(ok1711)* IV (3x outcrossed strain VC1233)

BJH2763: *ocr-2(ak47)* IV (3x outcrossed strain VM396)

BJH2764: *grk-2(gk268)* III *; flp-21(ok889)* V *; flp-18(db99)* X

NQ1298: *ceh-17(np1)* I *; grk-2(gk268)* III

XZ1557*: grk-2(gk268)* III ; *egl-4(n479)* IV

XZ1563: *grk-2(gk268)* III *; yakEx53[osm-6p::grk-2 cDNA::tbb-2* 3'UTR::OPERON::GFP*, myo-3p::mCherry]*

XZ1569: *grk-2(gk268)* III *pdfr-1(ok3425)* III

XZ1641: *grk-2(gk268)* III ; *yakEx71[xbx-1p::grk-2 cDNA::tbb-2* 3'UTR::OPERON::GFP*, myo-2p::mCherry]*

XZ2195: *yakSi32[nmr-1p::grk-2 cDNA::tbb-2* 3'UTR::OPERON::GFP*, cb-unc-119(+)]* II ; *grk-2(gk268)* III

XZ2241: *grk-2(gk268)* III ; *che-2(e1033)* X

XZ2242: *aptf-1(gk794)* II ; *grk-2(gk268)* III

XZ2243*: grk-2(gk268)* III ; *flp-11(tm2706)* X

XZ2244*: grk-2(gk268)* III ; *flp-18(db99)* X

XZ2245: *tph-1(mg280)* II ; *grk-2(gk268)* III

XZ2246: *grk-2(gk268)* III ; *yakEx189[odr-3p::grk-2 cDNA::tbb-2* 3'UTR::OPERON::GFP*, myo-2p::mCherry]*

XZ2249: *grk-2(gk268)* III ; *yakEx191[sra-6p::grk-2 cDNA::tbb-2* 3'UTR::OPERON::GFP, *myo-2p::mCherry]*

XZ2252: *grk-2(gk268)* III ; *yakEx194[str-1p::grk-2cDNA, str-2p::grk-2 cDNA, odr-10p::grk-2 cDNA, myo-2p::mCherry]*

XZ2253: *grk-2(gk268)* III ; *kin-29(oy38)* X

XZ2273: *grk-2(gk268)* III ; *yakEx202[gcy-8p::grk-2 cDNA::tbb-2* 3'UTR::Operon::GFP, *myo-2p::mCherry]*

XZ2277: *grk-2(gk268)* III ; *npr-1(ky13)* X *; yakEx206[ncs-1p::CRE, flp-21p::loxP::STOP::loxP::npr-1 cDNA::SL2::GFP, myo-2p::mCherry]*

XZ2278: *grk-2(gk268)* III ; *npr-1(ky13)* X

XZ2285: *grk-2(gk268)* III ; *yakIs19[GRK-2::tagRFP]; yakEx204[odr-3p*::mNeon::NLS]

XZ2291: *grk-2(gk268)* III ; *npr-1(ky13) X ; yakEx209[sra-6p::npr-1 cDNA, myo-2p*::mCherry*]*

XZ2309: *grk-2(gk268)* III ; *flp-1(ok2811) IV; flp-18(db99)* X

XZ2314: *flp-1(ok2811)* IV

XZ2317: *grk-2(gk268)* III ; *flp-1(ok2811)* IV ; *yakEx216*[*twk-47p::flp-1* cDNA::*tbb-2* 3'UTR::OPERON::GFP, *unc-122*::GFP]

XZ2345: *grk-2(gk268)* III *; egl-4(n479)* IV ; *yakEx221[osm-6p::egl-4 cDNA::tbb-2* 3'UTR::OPERON::GFP*, unc-122*::GFP]

XZ2379: *grk-2(gk268)* III ; *flp-1(ok2811)* IV ; *yakEx231*[*flp-1p*::FLP-1 ORF::GFP, *rab-3p*::mCherry]

XZ2404: *grk-2(gk268)* III ; *frpr-7(gk463846)* X

XZ2405: *grk-2(gk268)* III ; *npr-6(tm1497)* X

XZ2482: *grk-2(gk268)* III ; *yakEx253*[*hsp-16.41p*::*grk-2* cDNA::*tbb-2* 3'UTR::OPERON::GFP, *unc-122*::GFP]

XZ2487: *flp-1(ok2811)* IV ; *yakEx256*[*flp-1p(trc)*::FLP-1::mCherry, *unc-122*::GFP]

XZ2488: *grk-2(gk268)* III ; *flp-1(ok2811)* IV ; *yakEx256*[*flp-1p(trc)*::FLP-1::mCherry, *unc-122*::GFP]

XZ2501: *grk-2(gk268)* III ; *yakEx259*[*dat-1p*::*grk-2* cDNA::*tbb-2* 3'UTR::OPERON::GFP, *unc-122*::GFP]

XZ2522: *yakEx261*[*flp-1p*::*flp-1(trc)::*FLP-1::mCherry, *unc-122*::: GFP]

XZ2555: *grk-2(gk268)* III ; *flp-1(ok2811)* IV ; *npr-1(ky13)* X

**S2 Table. List of plasmids**

<u>Gateway destination vectors</u>

| | |
|---|---|
| pCFJ150 | Gateway destination vector for insertion at chr II Mos site *ttTi5605* |

<u>Gateway entry clones</u>

| | |
|---|---|
| BJP-T11 | *dat-1p* [4-1] (690 bp of the *dat-1* promoter upstream of the ATG) |
| BJP-C664 | *twk-47p* [4-1] (222 bp of the *twk-47* promoter upstream of the ATG) |
| pADA126 | *let-858* 3'UTR [2-3] |
| pCFJ31 | *acr-2p* [4-1] (3362 bp of the *acr-2* promoter upstream of the ATG) |
| pCFJ326 | *tbb-2* 3'UTR::OPERON::GFP [2-3] |
| pCFJ1973 | mNEON-NLS [1-2] |
| pCR185 | GFP::*unc-54* 3'UTR [2-3] |
| pEGB05 | *rab-3p* [4-1] (1224 bp of the *rab-3* promoter upstream of the ATG) |
| pET68 | *grk-2* cDNA [1-2] |
| pET85 | *osm-6p* [4-1] (2400 bp of the *nmr-1* promoter upstream of the ATG) |
| pET89 | *grk-2p* [4-1] (2895 bp of the *grk-2* promoter upstream of the ATG) |
| pET108 | *xbx-1p* [4-1] (425 bp of the *xbx-1* promoter upstream of the ATG) |
| pET276 | *odr-3p* [4-1] (4125 bp of the *odr-3* promoter upstream of the ATG) |

pET299 *gcy-8p* [4-1] (1923 bp of the *gcy-8* promoter upstream of the ATG)

pET312 *flp-1* cDNA [1-2]

pET332 *flp-1p* [4-1] (514 bp of the *flp-1* promoter upstream of the ATG)

pET336 *flp-1* ORF [1-2]

pGH107 tagRFP::*let-858* 3'UTR [2-3]

pIR47 *str-1p* [4-1] (4012 bp of the *str-1* promoter upstream of the ATG)

pIR211 *str-2p* [4-1] (2000 bp of the *str-2* promoter upstream of the ATG)

pIR419 *odr-10p* [4-1] (1000 bp of the *odr-10* promoter upstream of the ATG)

pJB-GL24 *sra-6p* [4-1] (2963 bp of the *sra-6* promoter upstream of the ATG)

pMA102 *nmr-1p* [4-1] (4709 bp of the *nmr-1* promoter upstream of the ATG)

<u>Gateway expression constructs</u>

pET79 *rab-3p::grk-2 cDNA:tbb-2* 3'UTR::OPERON::GFP_pCFJ150

pET83 *acr-2p::grk-2 cDNA::tbb-2* 3'UTR::OPERON::GFP_pCFJ150

pET86 *osm-6p::grk-2 cDNA::tbb-2* 3'UTR::OPERON::GFP_pCFJ150

pET90 *grk-2p::grk-2 cDNA*::GFP_pCFJ150

pET91 *grk-2p::grk-2 cDNA*::tagRFP_pCFJ150

pET109 *xbx-1p::grk-2 cDNA::tbb-2* 3'UTR::OPERON::GFP_pCFJ150

pET119 *nmr-1p::grk-2 cDNA::tbb-2* 3'UTR::OPERON::GFP_pCFJ150

pET278 *odr-3p::grk-2 cDNA::tbb-2* 3'UTR::OPERON::GFP_pCFJ150

pET285 *sra-6p::grk-2 cDNA::tbb-2* 3'UTR::OPERON::GFP_pCFJ150

pET286 *odr-10p::grk-2 cDNA::tbb-2* 3'UTR::OPERON::GFP_pCFJ150

pET289 *str-1p::grk-2 cDNA::tbb-2* 3'UTR::OPERON::GFP_pCFJ150

pET291 *str-2p::grk-2 cDNA::tbb-2* 3'UTR::OPERON::GFP_pCFJ150

pET304 *gcy-8p::grk-2 cDNA::tbb-2* 3'UTR::OPERON::GFP_pCFJ150

pET307 *odr-3p::mNeon::NLS::let-858* 3'UTR_pCFJ150

| | |
|---|---|
| pET308 | *sra-6p::npr-1 cDNA::let-858* 3'UTR_pCFJ150 |
| pET320 | *twk-47p*::*flp-1* cDNA::*tbb-2* 3'UTR::OPERON::GFP_pCFJ150 |
| pET326 | *osm-6p::egl-4 cDNA::tbb-2* 3'UTR::OPERON::GFP_pCFJ150 |
| pET344 | *flp-1p*::*flp-1* cDNA::*tbb-2* 3'UTR::OPERON::GFP_pCFJ150 |
| pET346 | *flp-1p*::FLP-1 ORF::GFP_pCFJ150 |
| pET362 | *hsp-16.41p::grk-2* cDNA::*tbb-2* 3'UTR::OPERON::GFP_pCFJ150 |
| pET370 | *dat-1p*::*grk-2* cDNA::*tbb-2* 3'UTR::OPERON::GFP_pCFJ150 |

<u>Gifts</u>

| | |
|---|---|
| pEM01 | *flp-21p::loxP::STOP::loxP::npr-1 cDNA::SL2::GFP* (a gift from Cori Bargmann) |
| pEM03 | *ncs-1p*::nCre (a gift from Cori Bargmann) |
| pCS232 | *flp-1(trc)p::*FLP-1::mCherry (a gift from Alexander Gottschalk) |
